# Physiologically based modeling of the effect of physiological and anthropometric variability on indocyanine green based liver function tests

**DOI:** 10.1101/2021.08.11.455999

**Authors:** Adrian Köller, Jan Grzegorzewski, Matthias König

## Abstract

Accurate evaluation of liver function is a central task in hepatology. Dynamic liver function tests (DLFT) based on the time-dependent elimination of a test substance provide an important tool for such a functional assessment. These tests are used in the diagnosis and monitoring of liver disease as well as in the planning of hepatobiliary surgery. A key challenge in the evaluation of liver function with DLFTs is the large inter-individual variability. Indocyanine green (ICG) is a widely applied test compound used for the evaluation of liver function. After an intravenous administration, pharmacokinetic (PK) parameters are calculated from the plasma disappearance curve of ICG which provide an estimate of liver function. The hepatic elimination of ICG is affected by physiological factors such as hepatic blood flow or binding of ICG to plasma proteins, anthropometric factors such as body weight, age, and sex, or the protein amount of the organic anion-transporting polypeptide 1B3 (OATP1B3) mediating the hepatic uptake of ICG. Being able to account for and better understand these various sources of inter-individual variability would allow to improve the power of ICG based DLFTs and move towards an individualized evaluation of liver function. Within this work we systematically analyzed the effect of various factors on ICG elimination by the means of computational modeling. For the analysis, a recently developed and validated physiologically based pharmacokinetics (PBPK) model of ICG distribution and hepatic elimination was utilized. Key results are (i) a systematic analysis of the variability in ICG elimination due to hepatic blood flow, cardiac output, OATP1B3 abundance, liver volume, body weight and plasma bilirubin level; (ii) the evaluation of the inter-individual variability in ICG elimination via a large *in silico* cohort of n=100000 subjects based on the NHANES cohort with special focus on stratification by age, sex, and body weight; (iii) the evaluation of the effect of various degrees of cirrhosis on variability in ICG elimination. The presented results are an important step towards individualizing liver function tests by elucidating the effects of confounding physiological and anthropometric parameters in the evaluation of liver function via ICG.

## 1 INTRODUCTION

Accurate evaluation of liver function is a central task in hepatology. Dynamic liver function tests (DLFT) based on time-dependent elimination of test substances provide an important tool for the functional assessment of the liver. These tests are used in the diagnosis and monitoring of liver disease as well as in the planning of hepatobiliary surgery.

Indocyanine green (ICG) is a widely applied test compound used for the evaluation of liver function. It is bound to plasma proteins in the blood, eliminated exclusively by the liver, and subsequently excreted into the bile. It is not reabsorbed by the intestinal tissue and therefore does not undergo enterohepatic circulation (Wheeler et al., 1958). After intravenous administration of ICG, the plasma disappearance curve of ICG can be measured either by repeated plasma sampling or through non-invasive fingertip methods. From the concentration-time profile a set of pharmacokinetic parameters can be calculated (e.g. ICG clearance, plasma disappearance rate (ICG-PDR), retention ratio after 15 minutes (ICG-R15), ICG half-life), which provide estimates of liver function based on the ICG elimination capacity of the liver (Köller et al., 2021; Sakka, 2018).

A major challenge in DLFTs based on the elimination of test substances such as ICG is the large inter-individual variability in their elimination. Physiological factors such as hepatic blood flow, binding to plasma proteins, or the amount of transport protein mediating the hepatic ICG uptake can influence the elimination of ICG. Furthermore, ICG elimination is generally reduced in liver disease and altered by surgical interventions such as partial hepatectomy (Köller et al., 2021).

### 1.1 Blood flow

The elimination of ICG depends strongly on hepatic blood flow due to its high hepatic extraction-ratio (0.6 - 0.9 in healthy subjects (Grainger et al., 1983)). The effect of varying hepatic blood flow on ICG-elimination has been studied by various groups. Kanstrup and Winkler found a strong positive correlation between hepatic blood flow and ICG-clearance when altering hepatic blood flow either pharmacologically or by food ingestion (Kanstrup and Winkler, 1987). Rowell et al. reported a strong positive correlation between changes in hepatic blood flow and ICG-clearance during exercise in healthy subjects but poor correlation in cirrhosis (Rowell et al., 1964). Gadano et al. and Huet and Villeneuve studied the dependency of ICG-clearance on hepatic blood flow in cirrhotic and healthy subjects (Gadano et al., 1997; Huet and Villeneuve, 1983). Huet and Lelorier were interested in the effects of smoking and chronic hepatitis B reporting hepatic blood flow and ICG parameters (Huet and Lelorier, 1980).

Hepatic blood flow results from the interplay of various factors such as blood pressure, portal resistance and cardiac output. Further, it is often altered in disease (e.g. in portal hypertension (Iwakiri, 2014)) and after liver surgery (e.g. partial hepatectomy (Kawasaki et al., 1991)). As a consequence, accurate evaluation of liver function via ICG pharmacokinetics is challenging in diseases that affect systemic circulation (e.g. cardiac output) or hepatic blood supply. A better understanding of the effects of blood flow on ICG elimination would enable a more accurate evaluation of ICG liver function tests.

### 1.2 Transport protein (OATP1B3)

ICG is removed from the blood via transporter-mediated uptake into the hepatocytes. Two proteins on the sinusoidal membrane are mainly responsible for the hepatic uptake of ICG in humans, the organic anion transporting polypeptide 1B3 (OATP1B3, gene symbol: SLCO1B3) and the Na^+^-taurocholate cotransporting polypeptide (NTCP) (de Graaf et al., 2011; Kagawa et al., 2017). OATP1B3 is the major transporter for hepatic ICG uptake and subjects with a homozygous SLCO1B3 null allele show markedly impaired ICG clearance (Anzai et al., 2020).

The OATP level in hepatocytes has a large impact on plasma concentrations of various drugs (Giacomini et al., 2010; Schipani et al., 2012; Schneck et al., 2004). In human subjects, a large variability in the protein amount of OATP1B3 exists (Burt et al., 2016; Kimoto et al., 2012; Prasad et al., 2014). The OATP1B3 level does not depend on age or sex (Prasad et al., 2014) though some dependency on ethnicity has been reported (Peng et al., 2015).

### 1.3 Plasma proteins and bilirubin

ICG is bound to plasma proteins (serum albumin and lipoproteins) (Kamisaka et al., 1974; Ott, 1998). It has been suggested that these plasma proteins are involved in the hepatic uptake mechanism of ICG at the sinusoidal membrane (Berk et al., 1987; Shinohara et al., 1996). However, no consensus about the effect of plasma proteins on ICG uptake has been reached. In contrast, the effect of bilirubin on ICG uptake has been extensively studied. Bilirubin is an end product of heme degradation which is primarily located in the blood and is almost completely bound to albumin (Fevery, 2008; Maruhashi et al., 2019). Significant negative correlation between ICG elimination and plasma bilirubin levels has been reported (Branch et al., 1976; Paumgartner et al., 1969). Changes in bilirubin plasma levels are often induced by hepato-biliary diseases, with the prime example being Gilbert’s disease (Fretzayas et al., 2012). Further, increased bilirubin plasma levels can be caused by biliary obstruction or viral hepatitis and are used as an indicator of acute liver failure (Sullivan and Rockey, 2017).

### 1.4 Anthropometric factors

An important question for the individualized and stratified evaluation via DLFTs is how factors such as body weight, age, and sex affect the elimination of the test substance. As summarized by Kim et al. (Kim et al., 2015), age has a significant effect on liver volume, blood flow, and function. A reduction of functional liver mass, total liver volume, and a decrease in hepatic blood flow are the most relevant age induced changes to the liver. Additionally, the susceptibility to liver disease increases with age accompanied by a decline in the hepatic regenerative response which has substantial consequences for liver surgery (Schmucker, 2005). Reduced ICG clearance with increasing age has been reported by Wood et al. (Wood et al., 1979). Liver volume and body weight have been shown to correlate well in western adults (Vauthey et al., 2002). Due to the dependency of ICG elimination on liver volume (Roberts et al., 1976), body weight is an important factor to be considered in the evaluation of liver function with ICG. Because of significant differences in average body weight and organ volumes between male and female subjects, sex is another important factor in the analysis of variability in ICG liver function tests. However, Martin et al. (Martin et al., 1975) reported no variations in ICG-clearance per kg body weight between male and female subjects, while reporting significantly increased ICG-PDR in women.

Within this work we systematically investigated the individual contribution of these factors to the large inter-individual variability in liver function tests based on ICG using a recently developed and validated physiologically based pharmacokinetics (PBPK) model of ICG (Köller et al., 2021).

## 2 MATERIAL AND METHODS

The presented work utilizes a recently established and validated PBPK model of ICG distribution and hepatic elimination (Fig. 1). The model is encoded in the Systems Biology Markup Language (SBML) (Hucka et al., 2019; Keating et al., 2020) and was developed using sbmlutils (König, 2021b), and cy3sbml (König and Rodriguez, 2019). Simulations were performed using sbmlsim (König, 2021a) based on the high-performance SBML simulator libroadrunner (Somogyi et al., 2015). We refer to (Köller et al., 2021) for details on the computational model, the model simulations, the data curation process, the calculation of pharmacokinetic parameters, and the implementation of liver cirrhosis. No model changes were performed compared to the model described in (Köller et al., 2021).

**Figure 1.**
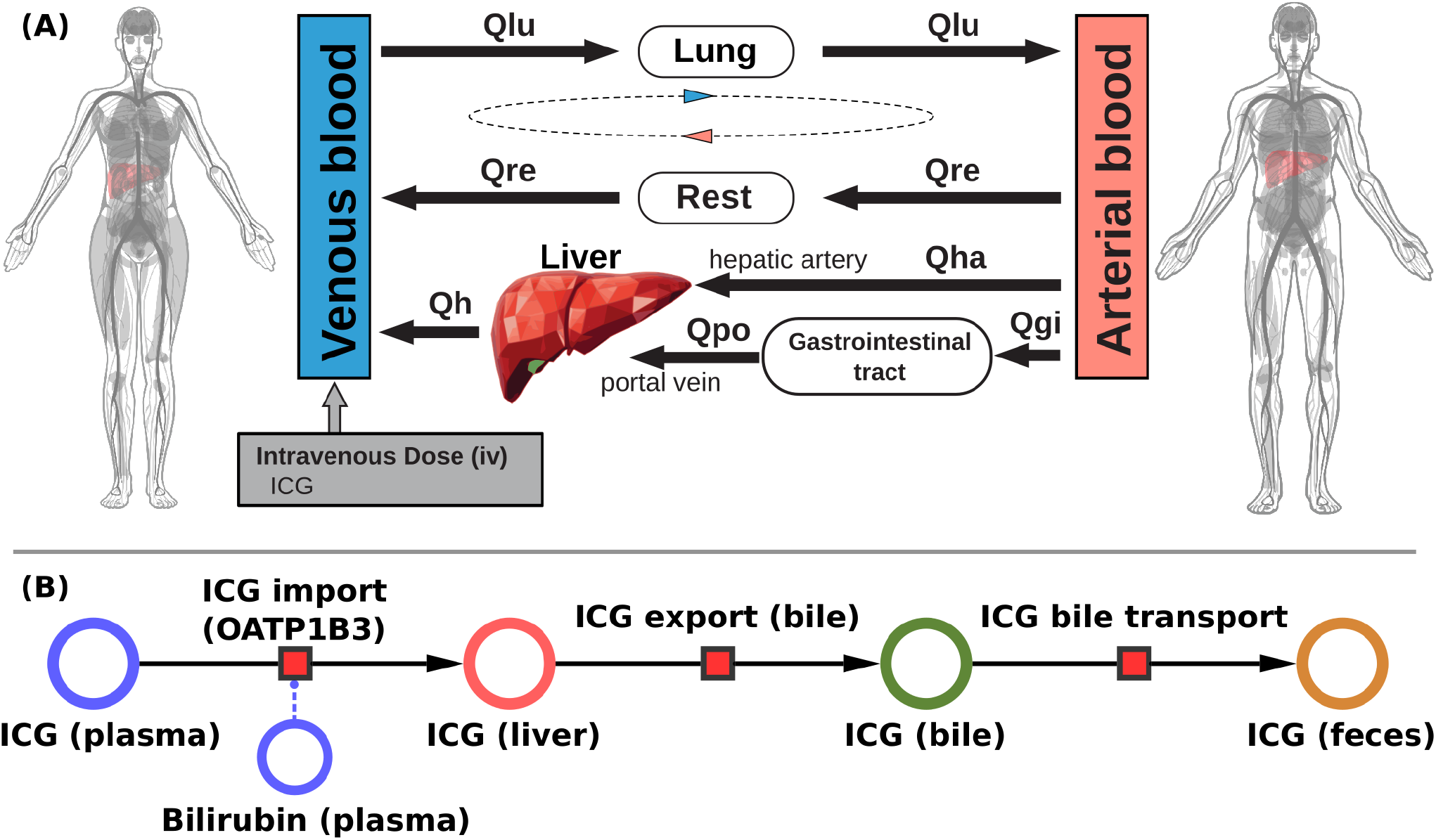
Model overview: **A:** PBPK model. The whole-body model for ICG consists of venous blood, arterial blood, lung, liver, gastrointestinal tract, and rest compartment (accounting for organs not modeled in detail). The systemic blood circulation connects these compartments. **B:** Liver model. ICG is taken up into the liver tissue (hepatocytes) via OATP1B3. The transport was modeled as competitively inhibited by plasma bilirubin. Hepatic ICG is excreted in the bile from where it is subsequently excreted in the feces. No metabolism of ICG occurs in the liver. Figure adapted from (Köller et al., 2021).

### 2.1 Parameter scans

Parameter scans (Fig. 2) were performed by varying the model parameters corresponding to the relative change in hepatic blood flow *f_bloodflow_* [-], relative change in cardiac output *f_cardiac_output_* [-], relative change in OATP1B3 amount *f*_*oatp*1*b*3_, liver volume *FVli* [l/kg], body weight *BW* [kg], and plasma bilirubin concentration *bil_ext_*[-]. The parameter values of the unchanged model are indicated as reference in the plots. For the relative changes *f_bloodflow_*, *f_cardiac_output_*, and *f*_*oatp*1*b*3_ the reference parameter value is 1.0 [-]. Bilirubin scans were performed on a log scale whereas all other scans were performed on a linear scale.

**Figure 2.**
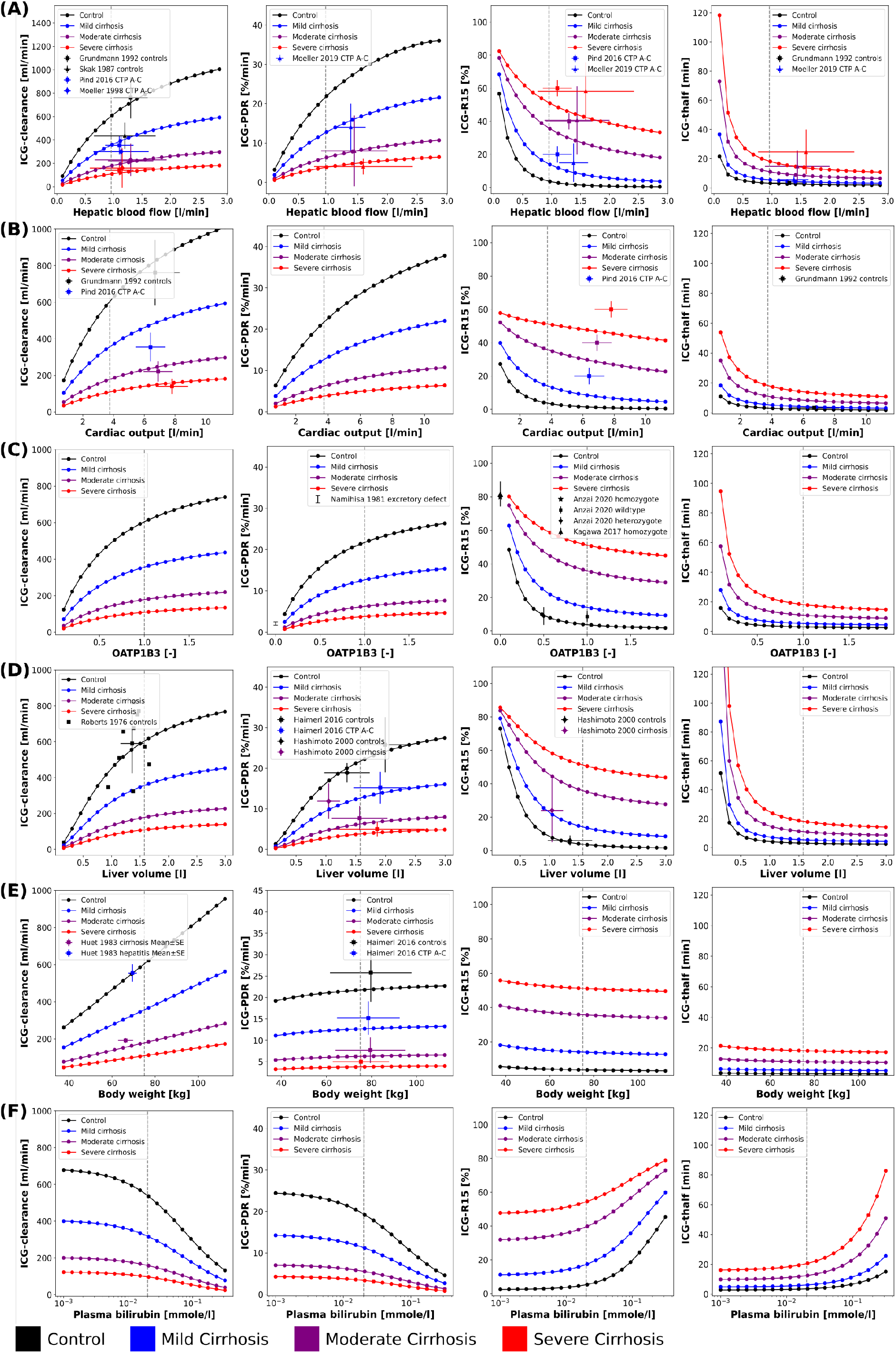
Effect of physiological factors on ICG parameters: The effect of hepatic blood flow, cardiac output, OATP1B3, liver volume, body weight, and plasma bilirubin on ICG-clearance, ICG-PDR, ICG-R15, and ICG-t_1/2_ were studied. Reference values of the model parameters, i.e. model parameters in the unchanged model, are indicated as vertical grey dashed lines. Black: control, blue: mild cirrhosis: purple: moderate cirrhosis, red: severe cirrhosis. **A:** Dependency on hepatic blood flow. Clinical data from Grundmann et al. (Grundmann et al., 1992), Pind et al. (Pind et al., 2016), Skak et al. (Skak and Keiding, 1987), and Moeller et al. (Møller et al., 1998, 2019). For additional data see Fig. 3. **B:** Dependency on cardiac output. Clinical data Grundmann et al. (Grundmann et al., 1992) and Pind et al. (Pind et al., 2016). **C:** Dependency on transport protein amount (OATP1B3). Clinical data from Anzai et al. (Anzai et al., 2020), Namihisa et al. (Namihisa et al., 1981), and Kagawa et al. (Kagawa et al., 2017). Heterozygote null variants with 0.5 activity, homozygote null variants with 0.0 activity. **D:** Dependency on liver volume. Clinical data from Haimerl et al. (Haimerl et al., 2016), Hashimoto et al. (Hashimoto and Watanabe, 2000), and Roberts et al. (Roberts et al., 1976). **E:** Dependency on body weight. Clinical data from Haimerl et al. (Haimerl et al., 2016) and Huet et al. (Huet and Villeneuve, 1983). **F:** Dependency on the plasma bilirubin concentration. For additional data see Fig. 4.

### 2.2 *In silico* population

For the analysis of variability in ICG pharmacokinetics, a large dataset of *in silico* individuals was created (see Fig. 5). Age, body weight, sex, height, and ethnicity were based on subjects sampled from the National Health and Nutrition Examination Survey (NHANES) spanning the years 1999-2018 (NHANES, 1999-2018). Due to the sampling from a real cohort it was possible to account for the covariances between the respective variables. Children and adolescents (age < 18 years), obese subjects (BMI > 30), and subjects with age ≥ 85 from 1999-2006 and ≥ 80 from 2007-2018 (due to top coding in NHANES at 85 and 80 years, respectively) were excluded. The resulting 35427 subjects were oversampled to reach 100000 individuals. This ensured sufficient samples in all stratified subgroups. Based on the age of a given individual, liver volume and hepatic blood flow were determined by multivariate sampling of liver volume per body weight and hepatic blood flow per body weight from the respective age distribution and subsequent multiplication with the individual body weight. This allowed to account for the covariances between liver volume and blood flow as well as the dependency of age. The normal distributions and covariance-matrices used for the sampling were calculated from data from Wynne et al. (Wynne et al., 1989). The transport protein amount (OATP1B3) was sampled independently from a lognormal distribution that was fitted to data from Peng et al. (Peng et al., 2015). The OATP1B3 distribution was assumed age and sex independent as reported by Prasad et al. (Prasad et al., 2014). Individual model simulations were performed by adjusting the model parameters corresponding to individual body weight *BW* [kg], liver volume *FVli* [l/kg], hepatic blood flow *f_bloodflow_* [-], and amount of OATP1B3 *f*_*oatp*1*b*3_ [-]. The scaling factors *f_bloodflow_* and *f*_*oatp*1*b*3_ describe the relative change from the model reference state. Individual ICG liver function test simulations were performed by administering 0.5 [mg/kg] ICG to the respective virtual individual and calculating the pharmacokinetics parameters on the resulting time course.

## 3 RESULTS

Within this work we systematically analyzed the effect of various factors on ICG elimination by the means of computational modeling. For the analysis, a recently developed and validated PBPK model of ICG distribution and hepatic elimination (see Fig. 1) was utilized (Köller et al., 2021). Model predictions were validated with data from multiple publications with an overview over all data sets provided in Tab. 1.

**Table 1.**
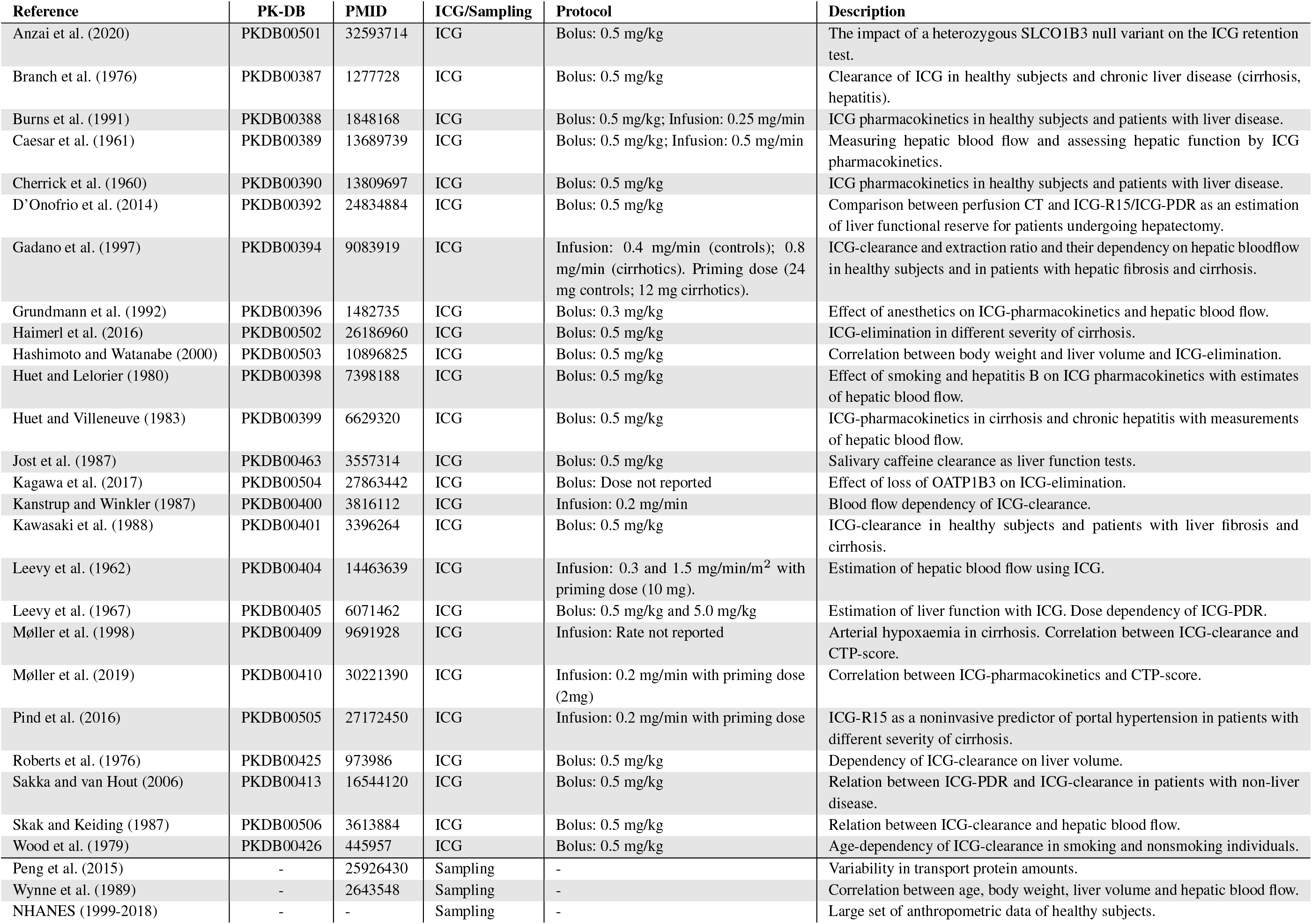
Overview of curated clinical studies.

### 3.1 PBPK Model

The systemic blood flow of the PBPK model is based on the cardiac output, which is scaled by the body weight. The liver is perfused through a dual blood supply consisting of arterial blood from the hepatic artery and venous blood from the portal vein. All organs besides the liver, the gastrointestinal tract, and the lung were pooled into the rest compartment. The included hepatic metabolism and biliary excretion are depicted in Fig. 1B. ICG is imported in the liver via OATP1B3. Hepatic ICG is exported through the bile and subsequently excreted via the feces. The effect of bilirubin on hepatic ICG uptake was modeled as a competitive inhibition based on the assumption that ICG can only be eliminated from the plasma if bound to plasma proteins. Due to the sequestration of plasma proteins bilirubin acts as a competitive inhibitor of ICG elimination, as both substances compete for plasma protein binding sites. Plasma bilirubin levels change slowly compared to the timescale of ICG liver function tests (15 min) and were therefore assumed in steady-state. Liver cirrhosis was modeled by a combination of functional tissue loss and shunts. For a detailed description of the model we refer to (Köller et al., 2021).

### 3.2 Variability due to physiological factors

First a systematic analysis of the effect of physiological variation on ICG parameters was performed (see Fig. 2). Specifically, the dependency of ICG-clearance, ICG-PDR, ICG-R15 and ICG-t_1/2_ on hepatic blood flow, cardiac output, transport protein amount (OATP1B3), liver volume, body weight, and plasma bilirubin was analyzed. An increase in all physiological factors except the plasma bilirubin concentration results in an increase in ICG elimination (increased ICG-PDR and ICG-clearance, reduced ICG-R15 and ICG-thalf). Changes in blood flow, OATP1B3 level, and liver volume have similar nonlinear effects on ICG parameters, whereas the body weight has an almost linear effect on ICG-clearance but only minimal effects on the other parameters. In the model, liver volume (tissue and blood vessels) are scaled with the body weight. As a result the plasma volume which is cleared from ICG every minute (clearance) increases with the body weight, without any change to the plasma disappearance rate, retention ratio or half-life.

Mild, moderate and severe cirrhosis reduce ICG elimination (decreased ICG-clearance and ICG-PDR, increased ICG-R15 and ICG-thalf) in a step-wise fashion without changing the overall curve shapes.

### 3.3 Validation of physiological factors

Model predictions were validated using various clinical data sets from healthy controls and subjects with mild, moderate and severe cirrhosis corresponding to Child-Turcotte-Pugh (CTP) classes A, B, and C, respectively (see Fig. 2: ICG-clearance, ICG-PDR, ICG-R15, and ICG-thalf depending on hepatic blood flow (Grundmann et al., 1992; Møller et al., 1998, 2019; Pind et al., 2016; Skak and Keiding, 1987); ICG-clearance, ICG-R15 and ICG-t_1/2_ depending on cardiac output (Grundmann et al., 1992; Pind et al., 2016); ICG-PDR and ICG-R15 depending on OATP1B3 (Anzai et al., 2020; Kagawa et al., 2017; Namihisa et al., 1981); ICG-clearance, ICG-PDR, and ICG-R15 depending on liver volume (Haimerl et al., 2016; Hashimoto and Watanabe, 2000; Roberts et al., 1976); ICG-clearance and ICG-PDR depending on body weight (Haimerl et al., 2016; Huet and Villeneuve, 1983). The model predictions show very good agreement with the validation data.

Additional validation of the model predictions were performed for the dependency of ICG parameters on blood flow (see Fig. 3) and plasma bilirubin (see Fig. 4).

**Figure 3.**
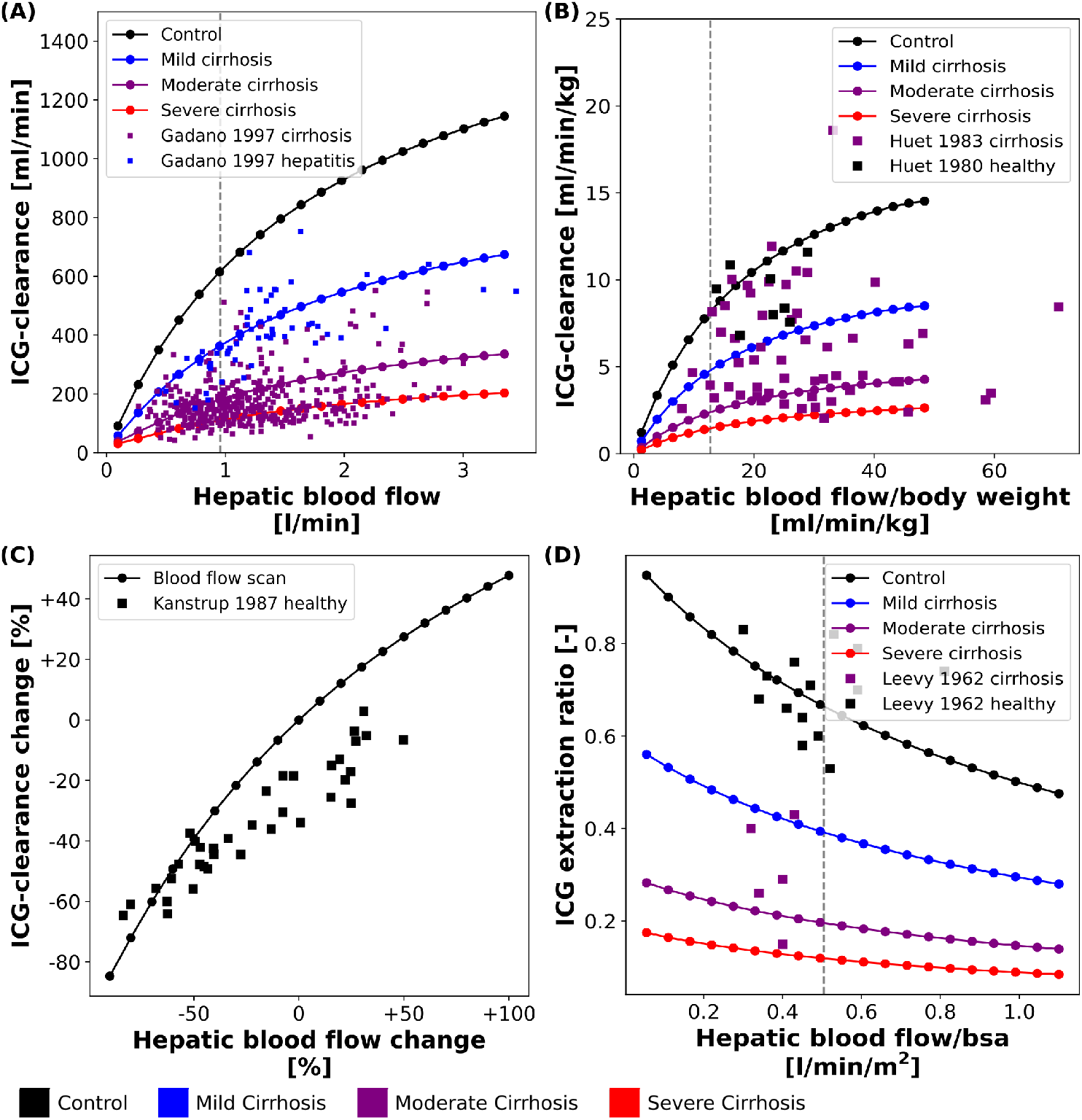
Effect of hepatic blood flow on ICG parameters: For model validation multiple blood flow experiments were simulated and compared to clinical data sets. Black: control, blue: mild cirrhosis: purple: moderate cirrhosis, red: severe cirrhosis. **A:** Dependency of ICG-clearance on hepatic blood flow. Clinical data of subjects with chronic hepatitis B and C and liver cirrhosis from Gadano et al. (Gadano et al., 1997). **B:** Dependency of ICG-clearance per kg body weight on hepatic blood flow per kg body weight. Clinical data of control and cirrhotic subjects from Huet et al. (Huet and Villeneuve, 1983; Huet and Lelorier, 1980). **C:** Dependency of ICG-clearance on externally changed hepatic blood flow. Clinical data of healthy subjects from Kanstrup and Winkler (Kanstrup and Winkler, 1987). **D:** Dependency of ICG extraction ratio on hepatic blood flow. Clinical data of control and cirrhotic subjects from Leevy et al. (Leevy et al., 1962).

**Figure 4.**
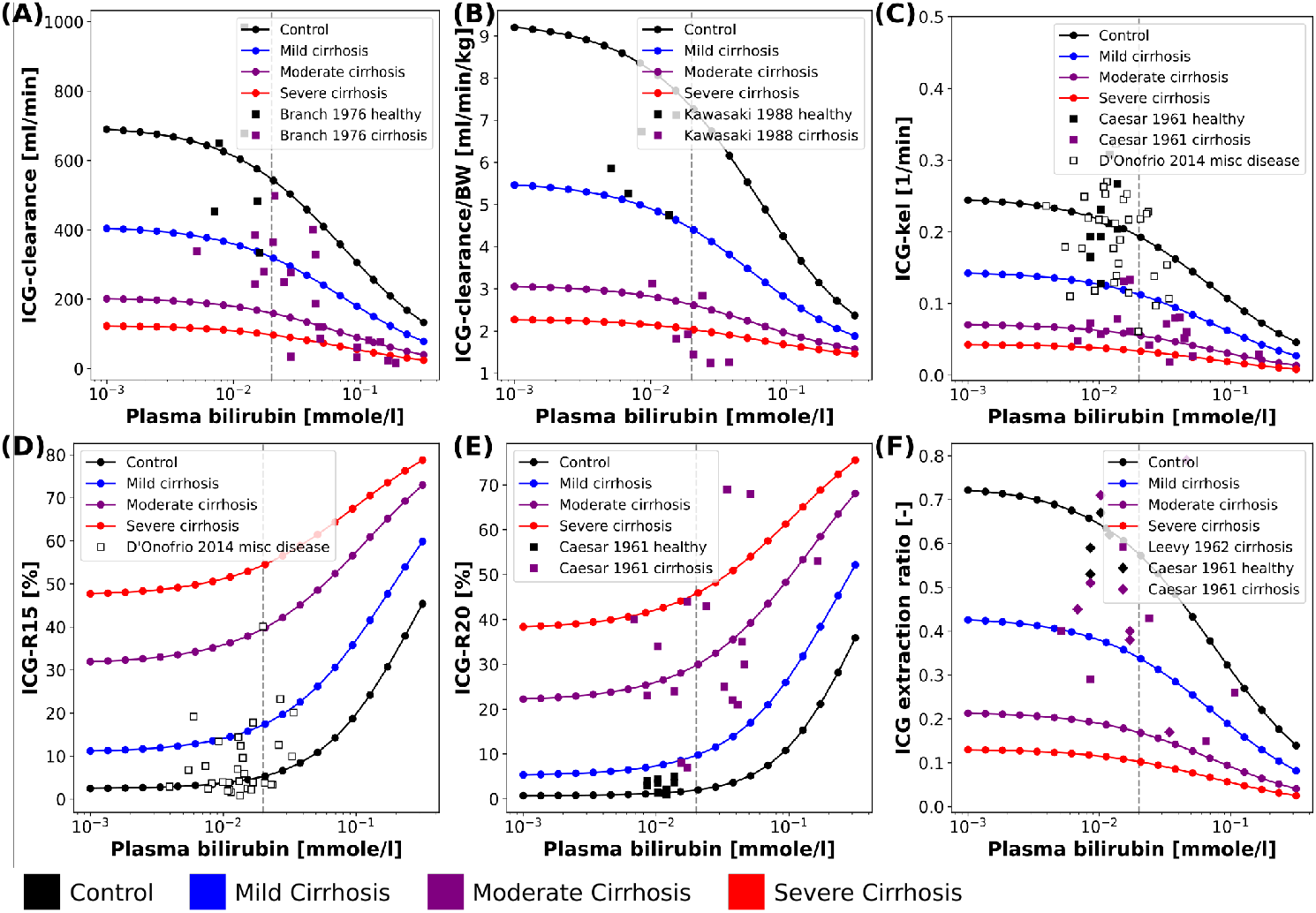
Effect of plasma bilirubin on ICG parameters: For model validation multiple plasma bilirubin experiments were simulated and compared to clinical data sets. Black: control, blue: mild cirrhosis: purple: moderate cirrhosis, red: severe cirrhosis. **A:** Dependency of ICG-clearance on the plasma bilirubin concentration. Clinical data of control and cirrhotic subjects from Branch et al. (Branch et al., 1976). **B:** Dependency of ICG-clearance per kg body weight on the plasma bilirubin concentration. Clinical data of control and cirrhotic subjects from Kawasaki et al. (Kawasaki et al., 1988). **C:** Dependency of ICG-kel on the plasma bilirubin concentration. Clinical data of control and cirrhotic subjects from Caesar et al. (Caesar et al., 1961). Clinical data of subjects with other liver diseases from D’Onofrio et al. (D’Onofrio et al., 2014). **D:** Dependency of ICG-R15 on the plasma bilirubin concentration. Clinical data of subjects with other liver diseases from D’Onofrio et al. (D’Onofrio et al., 2014). **E:** Dependency of ICG-R20 on the plasma bilirubin concentration. Clinical data of control and cirrhotic subjects from Caesar et al. (Caesar et al., 1961). **F:** Dependency of ICG extraction ratio on the plasma bilirubin concentration. Clinical data of control and cirrhotic subjects from Caesar et al. and Leevy et al. (Caesar et al., 1961; Leevy et al., 1962).

**Figure 5.**
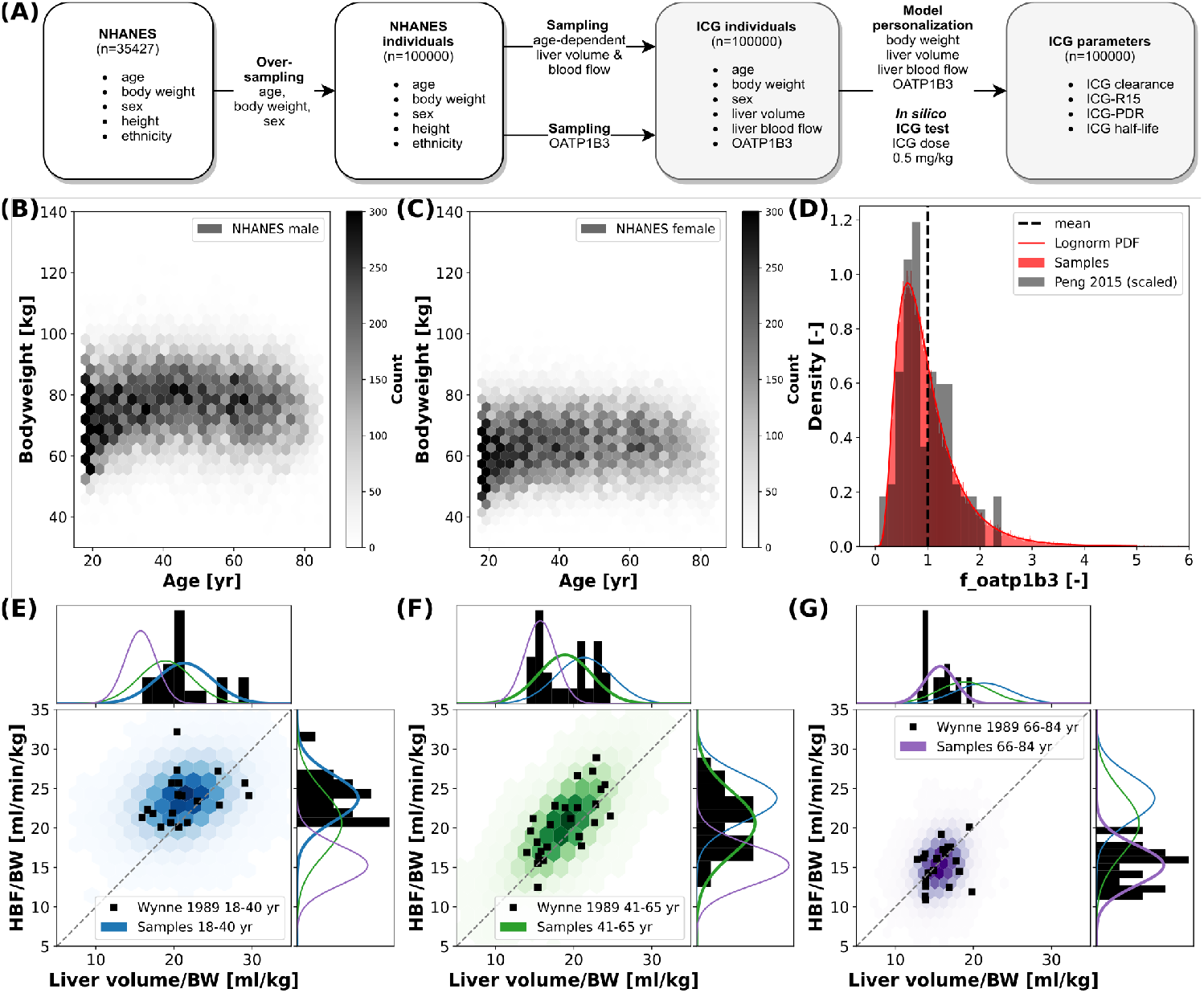
*In silico* population: ICG elimination was simulated for an *In silico* population consisting of n=100000 individuals (see Fig. 1E for workflow). Subjects are sampled form the NHANES cohort thereby accounting for the existing covariances between sex, age, body weight, height and ethnicity. **A:** Workflow to simulate ICG elimination in a large *in silico* population consisting of n=100000 individuals. **B, C:** Dependency of body weight on age in men and women. Data from NHANES (NHANES, 1999-2018) depicted as hexbin with shading corresponding to number of subjects (hexbins containing ≥ 300 subjects are encoded in black). **D:** Lognormal density distribution of OATP1B3 amount. Clinical data (grey) from Peng et al. (Peng et al., 2015) was normalized to the the model parameter f_oatp1b3 describing the change in OATP1B3 amount relative to the model reference value. **E-G:** Distributions of hepatic blood flow per kg body weight and liver volume per kg body weight in 3 different age groups: blue - 18-40 yr; green - 41-65 yr; purple - 66-84 yr. Clinical data (black) from Wynne et al. (Wynne et al., 1989).

Fig. 3A and B show the relation between ICG-clearance and hepatic blood flow in healthy subjects as well as patients of chronic hepatitis B or C and liver cirrhosis. ICG-clearance is reduced in liver disease, more severely in liver cirrhosis than hepatitis. The simulations were performed for healthy controls and three different degrees of cirrhosis (mild, moderate, severe) and are in good agreement with the clinical data (Gadano et al., 1997; Huet and Lelorier, 1980; Huet and Villeneuve, 1983). Subjects with hepatitis compare well to the simulation of moderate cirrhosis. Fig. 3C shows the change in ICG-clearance due to an externally induced change in hepatic blood flow by exercise, food ingestion or pharmacological intervention (Kanstrup and Winkler, 1987) with the corresponding model simulation. The relation is predicted accurately by the model, with an increase in blood flow resulting in an increased clearance of ICG. In Fig. 3D the dependency of the ICG extraction ratio on the hepatic blood flow is depicted. The extraction-ratio is the fraction of drug, that is removed from the plasma after a single pass through the liver. With increasing hepatic blood flow the extraction ratio decreases with model predictions for healthy and cirrhotic subjects being in good agreement with the data (Leevy et al., 1962).

The bilirubin plasma concentration in healthy subjects shows little variation (Fig. 4A-C, E, F). However, in liver cirrhosis and other liver diseases bilirubin levels may vary by an order of magnitude and are generally increased. The simulations depicted in Fig. 4 were performed for healthy controls and three different cirrhosis degrees based on the assumption of competitive inhibition of ICG uptake by bilirubin. Results were compared to clinical data of control and diseased subjects. Fig. 4 shows the dependency of ICG parameters on the plasma bilirubin levels (A, B: ICG-clearance; C: ICG-kel; D: ICG-R15; E: ICG-R20; F: ICG extraction ratio). Overall the decrease in ICG elimination due to elevated bilirubin plasma levels is predicted accurately. The simulations for healthy controls agree well with the data in healthy subjects.

In summary, model predictions for the effect of hepatic blood flow, cardiac output, OATP1B3 level, liver volume, body weight and bilirubin on ICG parameters could be validated with a large set of studies in healthy controls and various degrees of cirrhosis spanning almost 60 years of clinical research (Anzai et al., 2020; Branch et al., 1976; Caesar et al., 1961; D’Onofrio et al., 2014; Gadano et al., 1997; Grundmann et al., 1992; Haimerl et al., 2016; Hashimoto and Watanabe, 2000; Huet and Lelorier, 1980; Huet and Villeneuve, 1983; Kagawa et al., 2017; Kanstrup and Winkler, 1987; Kawasaki et al., 1988; Leevy et al., 1962, 1967; Møller et al., 1998, 2019; Namihisa et al., 1981; Pind et al., 2016; Roberts et al., 1976; Skak and Keiding, 1987).

### 3.4 *In silico* population

To analyze the inter-individual variability of ICG parameters, *in silico* ICG liver function tests were performed for 100000 subjects (Fig. 5. For every individual the age, body weight, sex, height and ethnicity was based on the NHANES cohort with subsequent sampling of age-dependent liver volume and hepatic blood flow and OATP1B3 amount. The 35427 NHANES subjects were oversampled to 100000 subjects, i.e., every subject occurred around three times with different liver volume and hepatic blood flow samples. By sampling from a large cohort of real individuals the covariances between the parameters age, body weight, sex, height and ethnicity could be accounted for. *In silico* ICG liver function tests used a dose of 0.5 [mg/kg] ICG (Fig. 5A). The dependency of the body weight on the age in men and women in the simulated population are depicted in Fig. 5B and C, respectively. In general, women have a lower body weight. Both distributions show an increase in body weight up until 40-60 year, followed by a decline in older ages. The number of subjects in younger age-groups is higher than in older age-groups, although this might not be representative of the population, but rather of the NHANES study design.

For every virtual individual the OATP1B3 level was sampled from a lognormal distribution based on reported protein concentrations by Peng et al. (Peng et al., 2015) (see Fig. 5D). OATP1B3 samples were assumed to be independent of age, body weight, sex, height and ethnicity.

For every virtual individual the age-dependent liver volume and hepatic blood flow were sampled under consideration of the underlying covariances between hepatic blood flow per kg body weight and liver volume per kg body weight, based on data by Wynne et al. (Wynne et al., 1989) (Fig. 5E-G). Both liver volume and hepatic blood flow per body weight decrease with increasing age. Age dependent multivariate sampling was used to determine hepatic blood flow and liver volume for every virtual individual, i.e. samples were taken from the respective age-dependent distribution.

### 3.5 Inter-individual variability

The inter-individual variability in ICG elimination was analyzed by performing an individualized ICG test simulation for each of the n=100000 subjects. ICG parameters were calculated from the individual ICG time courses. To analyse the effect of age, body weight and sex on ICG elimination the population was stratified by age (18-40 yr, 41-65 yr, 66-84 yr), body weight (40-60 kg, 60-80 kg, 80-100 kg, 100-140 kg), and sex (M, F). ICG clearance, PDR, R15, and t_1/2_ for each subgroup are shown in Fig. 6. With increasing age (independent of body weight), ICG elimination is reduced, as reflected by a decrease in clearance and PDR and an increase in ICG-R15 and t_1/2_. With increasing body weight ICG clearance increases whereas PDR, R15 and t_1/2_ show no change. Variation in body weight has a much larger effect on ICG-clearance, than on any other PK parameter in line with Fig. 2E.

**Figure 6.**
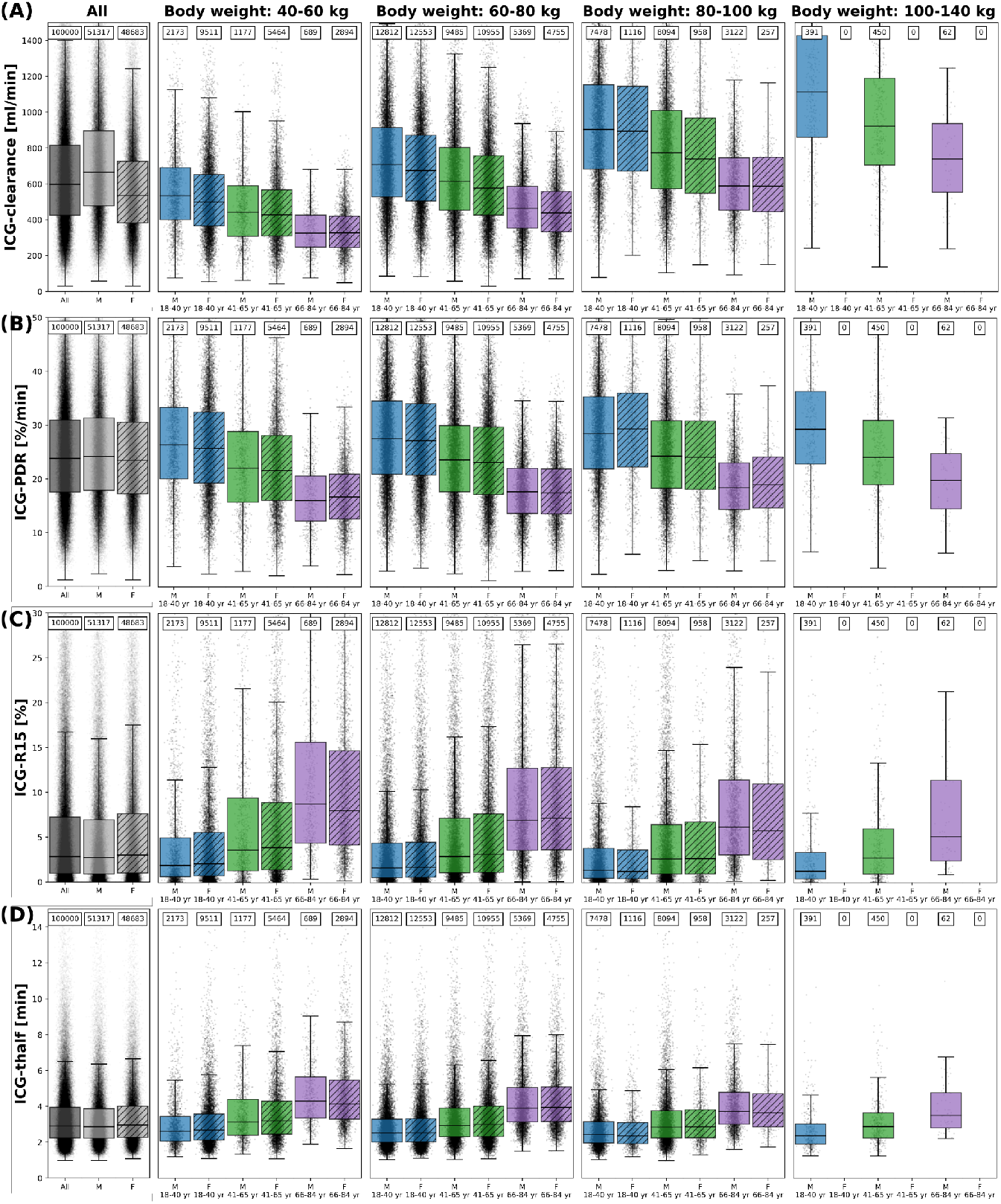
ICG parameters healthy *in silico* population: The dependency of **A** ICG-clearance [ml/min], **B** ICG-PDR [%/min], **C** ICG-R15[%], and **D** ICG-t_1/2_ [min] on body weight, sex and age in n=100000 individuals. Results are stratified by body weight class (40-60 kg, 60-80 kg, 80-100 kg, 100-140 kg), age group (blue - 18-40 yr; green - 41-65 yr; purple - 66-84 yr) and sex (M - unshaded; F - shaded). No women exist with body weight > 100 kg in the cohort. The sample size of the respective subgroups are depicted above each boxplot. The box extends from the lower to upper quartile values of the data, with a line at the median with whiskers as defined by Tukey. Individual data points are plotted for all subgroups. Subjects were simulated as healthy controls. For the corresponding results in mild cirrhosis, moderate cirrhosis, and severe cirrhosis see Supplementary Figure 1, 2 and 3, respectively.

Differences between the sexes are marginal in all parameters after stratification for body weight and age. Due to the lower body weight of women as depicted in Fig. 5B and C and the body weight dependency of ICG clearance an apparent difference between male and female ICG-clearance can be observed in the unstratified data.

The corresponding dependencies in mild cirrhosis, moderate cirrhosis, and severe cirrhosis are depicted in Supplementary Fig. 1, 2, and 3, respectively. Similar to healthy subjects, ICG elimination is reduced in cirrhosis with increasing age, as reflected by a decrease in clearance and PDR and an increase in ICG-R15 and t_1/2_. With increasing cirrhosis degree ICG clearance and PDR decrease and R15 and t_1/2_ increase, in line with Fig. 2.

### 3.6 Validation of population variability

To validate the results of the population model predictions (Fig. 6) multiple comparisons to clinical data sets were performed (Fig. 7 and Fig. 8).

**Figure 7.**
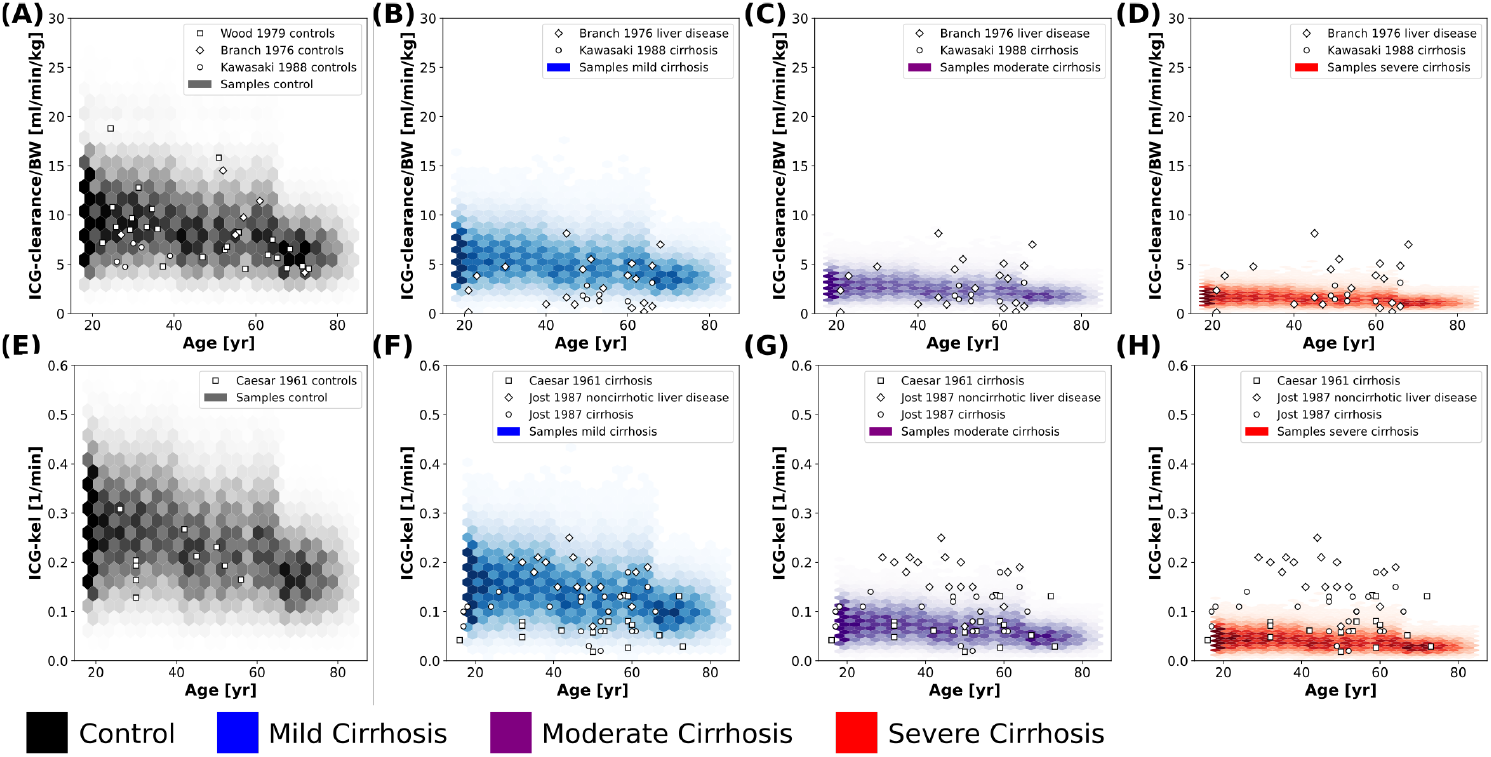
Validation of in *silico* population predictions: Age dependency of ICG clearance per body weight **A-D** and kel **E-H** in the population depicted as hexbins. Simulations were performed for healthy control (black), mild cirrhosis (blue), moderate cirrhosis (purple), and severe cirrhosis (red). Clinical data of controls from Wood et al., Branch et al., Kawasaki et al, and Caesar et al. (Wood et al., 1979; Branch et al., 1976; Kawasaki et al., 1988; Caesar et al., 1961) and cirrhotic subjects from Branch et al., Kawasaki et al., Caesar et al., and Jost et al. (Branch et al., 1976; Kawasaki et al., 1988; Caesar et al., 1961; Jost et al., 1987).

**Figure 8.**
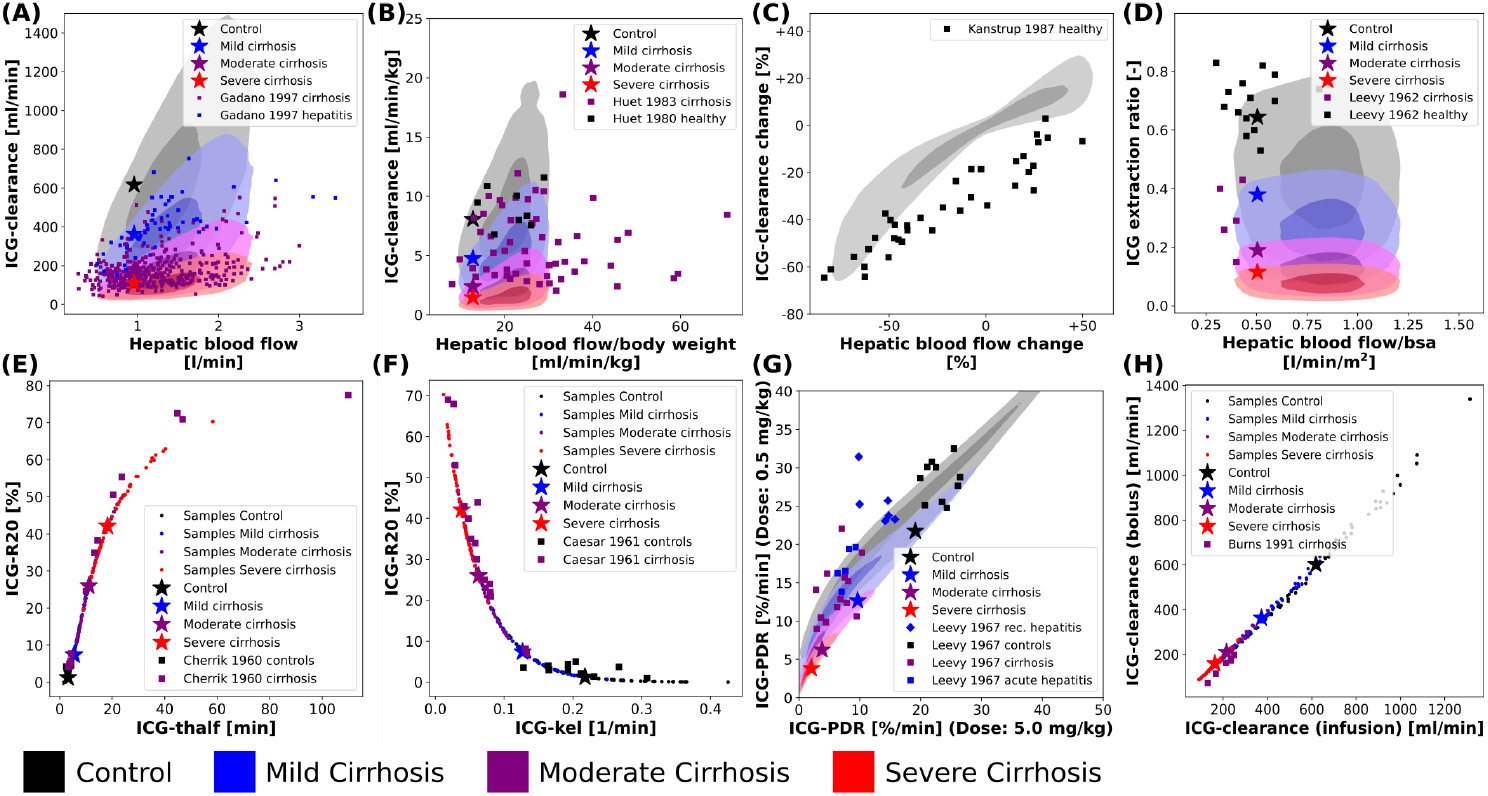
Variability analysis - model validation: The simulations were performed for four different cirrhosis degrees (except in **C**). Black: control, blue: mild cirrhosis: purple: moderate cirrhosis, red: severe cirrhosis. Stars depict the respective reference simulations. **A-D, G:** The identical simulations as in Fig. 3 were performed for the entire *in silico* population (n=100000). **E, F and H:** Simulations were performed for a subset of the *in silico* population (n=50). **E:** Correlation between ICG-R20 and ICG-tļ/2. Clinical data of control and cirrhotic subjects from Cherrick et al. (Cherrick et al., 1960). **F:** Correlation between ICG-R20 and ICG-kel. Clinical data of control and cirrhotic subjects from Caesar et al. (Caesar et al., 1961). **G:** Correlation between ICG-PDR after an ICG dose of 0.5 mg/kg and 5.0 mg/kg. Clinical data of control subjects and subjects with hepatitis (recovering and acute) and liver cirrhosis from Leevy et al. (Leevy et al., 1967). **H:** Correlation between ICG-clearance after a bolus administration and a constant infusion of ICG. Clinical data of cirrhotic subjects from Burns et al. (Burns et al., 1991)

The predicted dependency of ICG-clearance and ICG-kel on age in the in *silico* population in Fig. 7 is in very good agreement with data for healthy controls (Branch et al., 1976; Caesar et al., 1961; Kawasaki et al., 1988; Wood et al., 1979), and cirrhosis (Branch et al., 1976; Caesar et al., 1961; Jost et al., 1987; Kawasaki et al., 1988). ICG-clearance and ICG-kel decrease with increasing age and with increasing cirrhosis degree. The clinical data for cirrhosis did not provide any information on the disease severity such as CTP scores, so that cirrhosis data is plotted for all simulated severities. The model predictions cover the complete range of reported ICG-clearance and ICG-kel in cirrhotic subjects.

For additional validation, the population variability in the dependency of ICG parameters on hepatic blood flow was studied in Fig. 8A-D) (corresponding to Fig. 3).

Fig. 8A and B show the effect of inter-individual differences on the dependency of ICG-clearance on hepatic blood flow. The resulting variability especially in the control samples (grey area), is very large emphasizing the importance of including confounding factors in the evaluation of liver function.

Fig. 8C shows the change in ICG-clearance when hepatic blood flow is changed either by exercise, food ingestion or by pharmacological intervention. By including the inter-individual differences in the simulation, we were able to predict the variability in this dependency accurately. From the clinical data, we assume however, that the external influences, that were used to alter hepatic blood flow, had additional effects on ICG elimination, because even subjects with unchanged hepatic blood flow showed a decrease in clearance. It would be of high interest to analyze what these effects might be. This also shows, that recent food ingestion and long-term medication that affects blood flow are additional factors that have to be considered in liver function evaluation with ICG.

Fig. 8D shows the relation between the ICG extraction ratio and the hepatic blood flow. The changes in extraction ratio are in good agreement with the clinical data, but the simulations taking inter-individual into account cover a much larger range of blood flow per body surface area than the data.

Next, the inter-individual variability in the correlations between ICG parameters in healthy and cirrhotic subjects were simulated (Fig. 8E-H). The corresponding results without variability have been published previously (Köller et al., 2021).

Interestingly, all variability in the correlation between ICG-R20 and ICG-t_1_ /_2_ as well as ICG-R20 and ICG-kel lies on a nonlinear relationship (see Fig. 8E and F). Moreover, the clinical data follows the same nonlinear dependencies. Because all PK parameters of a subject are calculated from the same plasma disappearance curve, the results follow a strict nonlinear relationship.

In Fig. 8G the correlation between ICG-PDR after ICG doses of 0.5 mg/kg and 5.0 mg/kg is depicted. The relation is mostly linear, however some variation is observed, which agrees very well with the clinical data. This suggests a small dose-dependency of ICG-PDR depending on individual subject properties. In a previous analysis we could not find any dose-dependency of ICG-PDR for the model with reference parameters (Köller et al., 2021). A more detailed investigation into the factors causing the PDR dose dependency would be highly relevant.

A linear correlation, without deviation due to inter-individual differences, is observed between ICG-clearance calculated after a bolus administration and during a constant intravenous infusion of ICG (see Fig. 8H) with the clinical data following the same linear dependency.

### 3.7 Contribution of individual factors

Finally, we were interested in the contribution of individual factors to the observed population variability (Fig. 9 using a data set of ICG-clearance and ICG-PDR by Sakka and van Hout (Sakka and van Hout, 2006). The predicted variability between ICG-clearance and ICG-PDR in the in *silico* population is in very good agreement with the observed variability in the data (Fig. 9A). The combined effect of variation in hepatic blood flow, OATP1B3 amount, liver volume and body weight predicts the variability in the clinical data very accurately.

**Figure 9.**
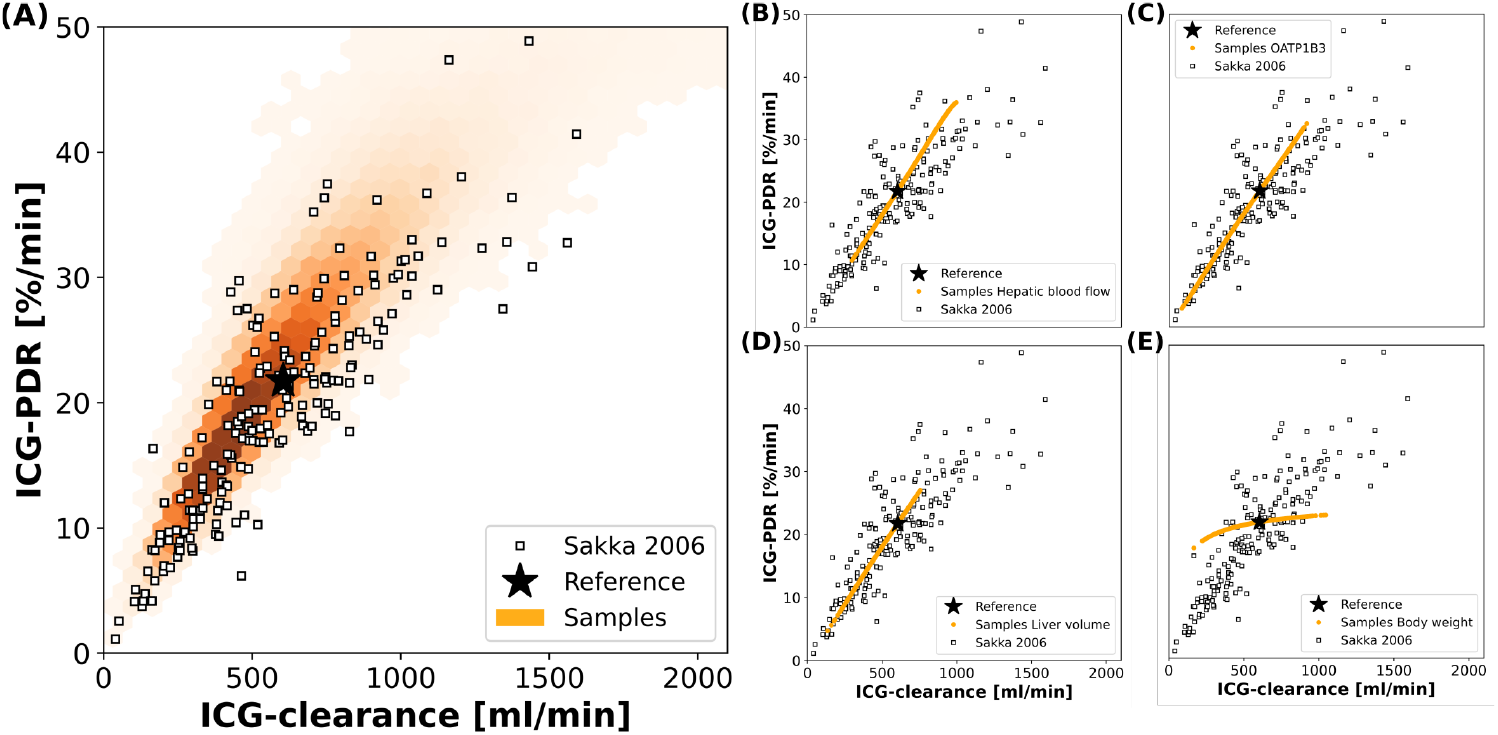
Contributions to population variability: Relationship between ICG-clearance and ICG-PDR with clinical data from Sakka and van Hout (Sakka and van Hout, 2006). **A:** Results of the simulations of the complete *in silico* population (n=100000). **B-E** Results of simulations, when hepatic blood flow, OATP1B3 level, liver volume, or body weight were varied individually in the range covered by the population. The respective other factors were set to their respective reference values in the simulation. Stars depict the reference simulation of the model.

To study the individual contribution of each of the above-mentioned factors these factors were varied individually while setting the remaining factors to their reference values (Fig. 9B-E). Hepatic blood flow, OATP1B3 amount, and liver volume affect ICG-clearance and PDR in a similar manner resulting in a linear dependency. Compared to the model’s reference simulation, these factors are able to increase as well as reduce ICG elimination. The individual effects appear to be additive as none of them individually can achieve the complete range of variability that is observed when they are varied together (Fig. 9A). In contrast, the body weight has a much smaller effect on ICG-PDR than on ICG-clearance thereby spanning an alternative axis of variation. Specifically, it is the main source of deviation from the correlation line between PDR and clearance.

## 4 DISCUSSION

Within this work, the effects of physiological and anthropometric factors on ICG elimination were studied systematically. For the analysis, a recently developed and validated PBPK model of ICG (see Fig. 1) was used (Köller et al., 2021).

The previous modeling approach was extended by the development of a large in *silico* population based on the NHANES cohort (NHANES, 1999-2018) with individualized model predictions for the n=100000 subjects. Our approach allowed to account for covariances between age, sex, body weight, height and ethnicity as well as age, liver volume and hepatic blood flow. Via individualized in *silico* ICG liver function tests, the inter-individual variability in ICG-elimination could be studied in healthy subjects as well as in mild, moderate, and severe cirrhosis corresponding to CTP-A, CTP-B, and CTP-C. Model predictions were validated with independent clinical data sets from 25 publications which were not used in the model development (see Tab. 1 and figures). Including the variability of physiological factors (hepatic blood flow, cardiac output, OATP1B3 level, liver volume, plasma bilirubin) and anthropometric parameters (body weight, age, sex) in the model simulations allowed to accurately predict the individual effects of these factors as well as the observed inter-individual variability in ICG elimination. The importance of including these factors in the evaluation of liver function with ICG is apparent.

Certain limitations exist in the presented analysis. First, the analysis focused specifically on factors for which (i) clinical data is available, (ii) alterations in ICG elimination are expected, (iii) and which could be included in our PBPK modeling approach. This analysis is far from exhaustive and other factors may exist which influence ICG-elimination but were not considered here. Second, the distributions of liver volume, hepatic blood flow and OATP1B3 amount were based on clinical data with relatively small sample size. The resulting distributions do not necessarily reflect the distribution and covariances within the NHANES population. For instance, some ethnical differences in OATP1B3 distribution may exist that were not considered here (see below for details). Furthermore, parameters which were assumed statistically independent could actually have dependencies. For example, an important assumption was that the amount of OATP1B3 is independent of other sampled factors such as body weight, age, sex, liver volume, and liver blood flow. Prasad et al. showed that OATP1B3 is independent of age and sex (Prasad et al., 2014), yet no data for the relationship between OATP1B3 amount and other anthropometric factors (body weight, liver volume, and liver blood flow) could be found in the literature.

A key factor for ICG elimination is the amount of OATP1B3 in the liver as it is the primary transport protein mediating hepatic ICG uptake. An increase in its amount results in increased ICG elimination (Fig-2C). A previous study has shown that ICG uptake into the liver correlates with the expression of OATP1B3 (Masuoka et al., 2020). Whereas some studies quantified the OATP1B3 amount in Human liver samples (Burt et al., 2016; Peng et al., 2015; Prasad et al., 2014) to our knowledge no data reporting OATP1B3 amount and systemic ICG parameters exists. For validation we compared reported ICG-PDR and ICG-R15 values for different genotypes. Under the assumed transport activities (wildtype enzyme 1.0, heterozygote null genotype 0.5, homozygote null genotype 0.0) for the model predictions, the simulations and experimental data are in very good agreement. Although OATP1B3 expression were significantly lower in the heterozygote than in the wild-type, the ICG-R15 results were comparable (Anzai et al., 2020), suggesting that some compensation is possible. Subjects with markedly poor ICG clearance but without severe liver disease are diagnosed with constitutional ICG excretory defect (Anzai et al., 2020; Namihisa et al., 1981) and lack of OATP1B3 expression has been confirmed in these subjects by immunohistochemistry (Kagawa et al., 2017; Masuoka et al., 2020). The incidence of ICG excretory defect is 0.007% in the Japanese population (Masuoka et al., 2020). Anzai et al. emphasized the importance of investigating the effect of the SLCO1B3 genotype on ICG clearance in cirrhotic patients in the future (Anzai et al., 2020). Within this work, this important investigation could be performed *in silico* (Fig. 2).

Whereas no differences in OATP1B3 expression with age or sex have been reported (Prasad et al., 2014), a difference in OATP1B3 amount with ethnicity may exist. Peng et al. showed based on a relatively small study (n=102 Caucasian, n=18 Asian, n=5 African-American) (Peng et al., 2015) that Asians have almost double the OATP1B3 amount with 27.6 [fmol/μg] (24.5 - 30.7 95% CI) of Caucasians with only 14.2 [fmol/μg] (12.9 - 15.6 95% CI). Ethnicity information was available for all our in *silico* subjects, but unfortunately not compatible with the ethnicity groups provided by Peng et al. E.g. Asians were not separately listed in NHANES but part of the larger group ’Other Race - Including Multi-Racial’. In summary, the available information on the ethnicity dependency of OATP1B3 expression was not sufficient to include it in our simulations.

The influence of hepatic blood flow on ICG-elimination was analyzed extensively (see Fig. 2A and B; Fig. 3; Fig. 8E-F). As can be observed in Fig 2A and B, the model’s reference values for cardiac output and hepatic blood flow consistently lie below the mean values from clinical data at the lower end of the physiological range. Using relatively low cardiac output in the reference state was necessary to accurately predict the ICG extraction ratio without implementing a longer delay between blood exiting the liver and entering the liver again (which would either require delayed differential equations or constructs such as the linear chain trick to achieve sufficient delays). This simplification had no effect on the ICG plasma disappearance curve and the resulting PK parameters. However, as a side effect of this, the hepatic blood flow of the reference simulations (stars) depicted in Fig. 8A, B and D are at the lower ends of the simulated range. This has no relevance for the variability analysis, but moving towards individualized ICG liver function tests the implementation of a circulation delay will be necessary.

In spite of the observed relationship between the plasma bilirubin concentration and ICG-elimination (see Fig. 2F and 4), it was not included in the population variability analysis. The main reason for this is the low variability of plasma bilirubin in healthy subjects compared to subjects with liver disease. In case of liver cirrhosis the observed relationship between bilirubin and ICG parameters is difficult to interpret. It is possible that the changes in ICG-elimination result from the disease itself and the observed changes in plasma bilirubin are simply a consequence of the liver disease without any effect on ICG uptake. Under the model assumption that bilirubin has a competitive inhibitory effect on ICG uptake the dependency of ICG parameters on bilirubin could be reproduced. In rat a clear effect of bilirubin plasma concentration (varied via injection) could be observed on ICG uptake, which supports our assumption (Paumgartner et al., 1969). Human data without liver disease but substantially altered bilirubin levels would be required to test the hypothesis that bilirubin is a competitive inhibitor of ICG uptake in humans.

Our study underlines the importance of quantifying and reporting factors affecting the inter-individual variability in liver function tests. The presented results have important implications for clinical testing of liver function with ICG as well as clinical studies involving ICG parameters. Most importantly, factors affecting ICG elimination must be quantified and reported alongside ICG parameters. Most of these factors can be easily determined in clinical practice such as body weight, sex, age in the anamnesis, plasma bilirubin via blood chemistry, hepatic blood flow and cardiac output via Doppler ultrasound, liver volume via imaging, and severity of cirrhosis using the CTP or MELD classification. The only factor difficult to obtain is the amount of OATP1B3 which would require invasive liver biopsy followed by protein quantification via immunohistochemistry. This could be a feasible approach in settings were liver biopsies are readily available, e.g., in hepatectomies. Clinical studies evaluating ICG elimination in different subgroups must account for differences in confounding factors such as age between groups. Especially in study designs comparing healthy controls (often young) with subjects with disease (often old) this is difficult to achieve. Our results provide quantitative information on the expected effect sizes.

In summary, the model shows a great potential in predicting the variability in ICG elimination in healthy as well as in cirrhotic subjects. By including variations in hepatic blood flow, hepatic transport proteins, liver volume, and body weight in the model simulation, new insights were gained on the correlation between ICG parameters and the dependency of the extraction ratio on hepatic blood flow. Important future questions are (i) how the information on underlying causes of inter-individual variability can be used for an improved evaluation of ICG based liver function tests; and (ii) how this variability influences risk assessment of postoperative survival after liver surgery based on ICG, e.g. the prediction of survival after partial hepatectomy as proposed in (Köller et al., 2021).

## AUTHOR CONTRIBUTIONS

AK and MK designed the study, developed the computational model, implemented and performed the analysis, and wrote the initial draft of the manuscript. JG provided support with PK-DB and data curation. All authors discussed the results. All authors contributed to and revised the manuscript critically.

## DATA AVAILABILITY STATEMENT

All clinical data of ICG pharmacokinetics that was used in this work can be found in PK-DB available from https://pk-db.com.

