## Supplementary Material for "Physiologically based modeling of the effect of physiological and anthropometric variability on indocyanine green based liver function tests"

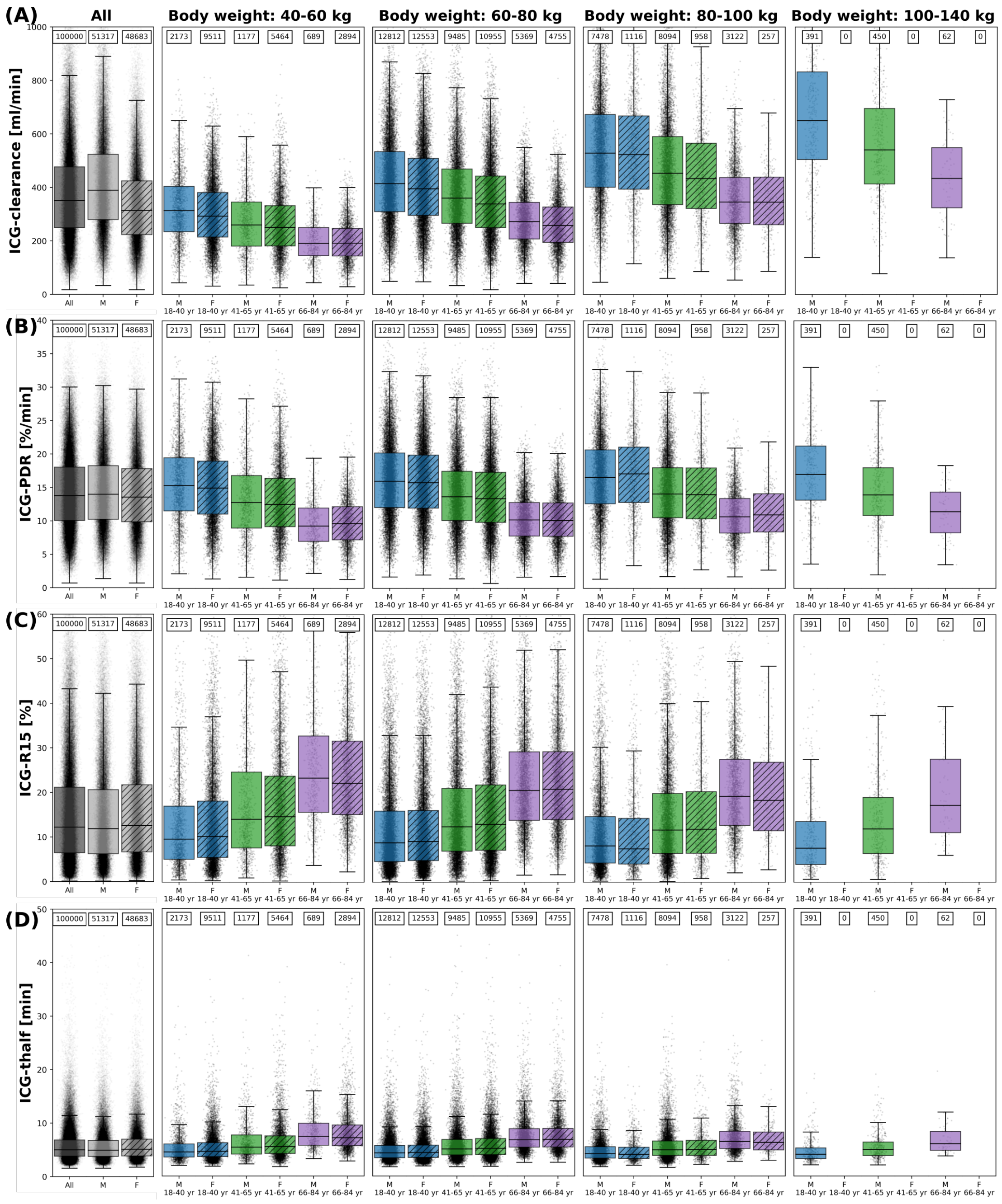


**Figure S1. ICG parameters in *in silico* population with mild cirrhosis:** The dependency of **A** ICG-clearance, **B** ICG-PDR, **C** ICG-R15, and **D** ICG-$t_{1/2}$ on body weight, sex and age in n=100000 individuals. Results are stratified by body weight class (40-60 kg, 60-80 kg, 80-100 kg, 100-140 kg), age group (blue - 18-40 yr; green - 41-65 yr; purple - 66-84 yr) and sex (M - unshaded; F - shaded). No women exist in the data set with body weight $>$ 100 kg. The sample size of the respective subgroups are depicted above each boxplot. The box extends from the lower to upper quartile values of the data, with a line at the median with whiskers as defined by Tukey. Individual data points are plotted for all subgroups. Subjects are simulated with mild cirrhosis.

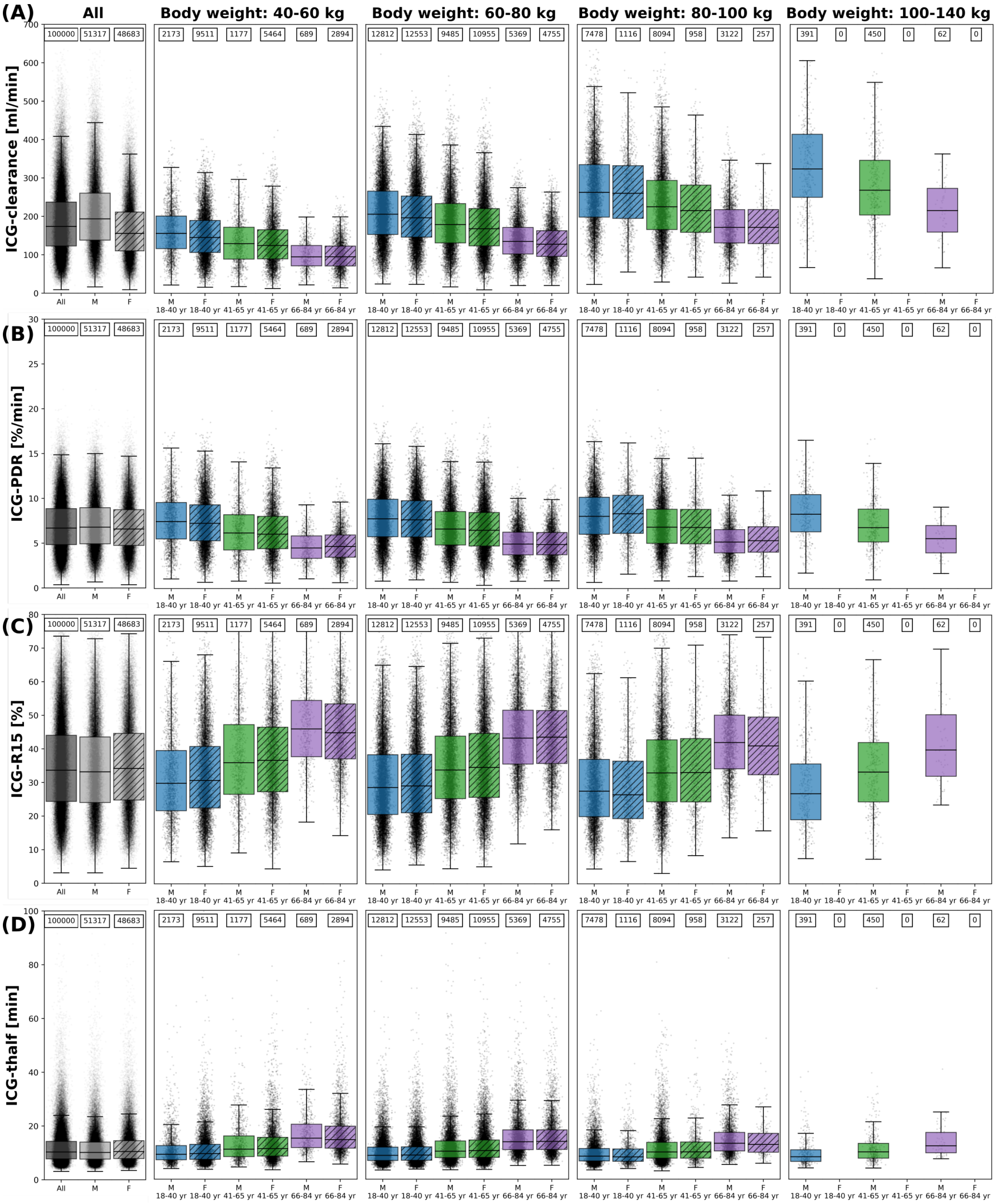


**Figure S2. ICG parameters in *in silico* population with moderate cirrhosis:** The dependency of **A** ICG-clearance, **B** ICG-PDR, **C** ICG-R15, and **D** ICG-$t_{1/2}$ on body weight, sex and age in n=100000 individuals. Results are stratified by body weight class (40-60 kg, 60-80 kg, 80-100 kg, 100-140 kg), age group (blue - 18-40 yr; green - 41-65 yr; purple - 66-84 yr) and sex (M - unshaded; F - shaded). No women exist in the data set with body weight > 100 kg. The sample size of the respective subgroups are depicted above each boxplot. The box extends from the lower to upper quartile values of the data, with a line at the median with whiskers as defined by Tukey. Individual data points are plotted for all subgroups. Subjects are simulated with moderate cirrhosis.

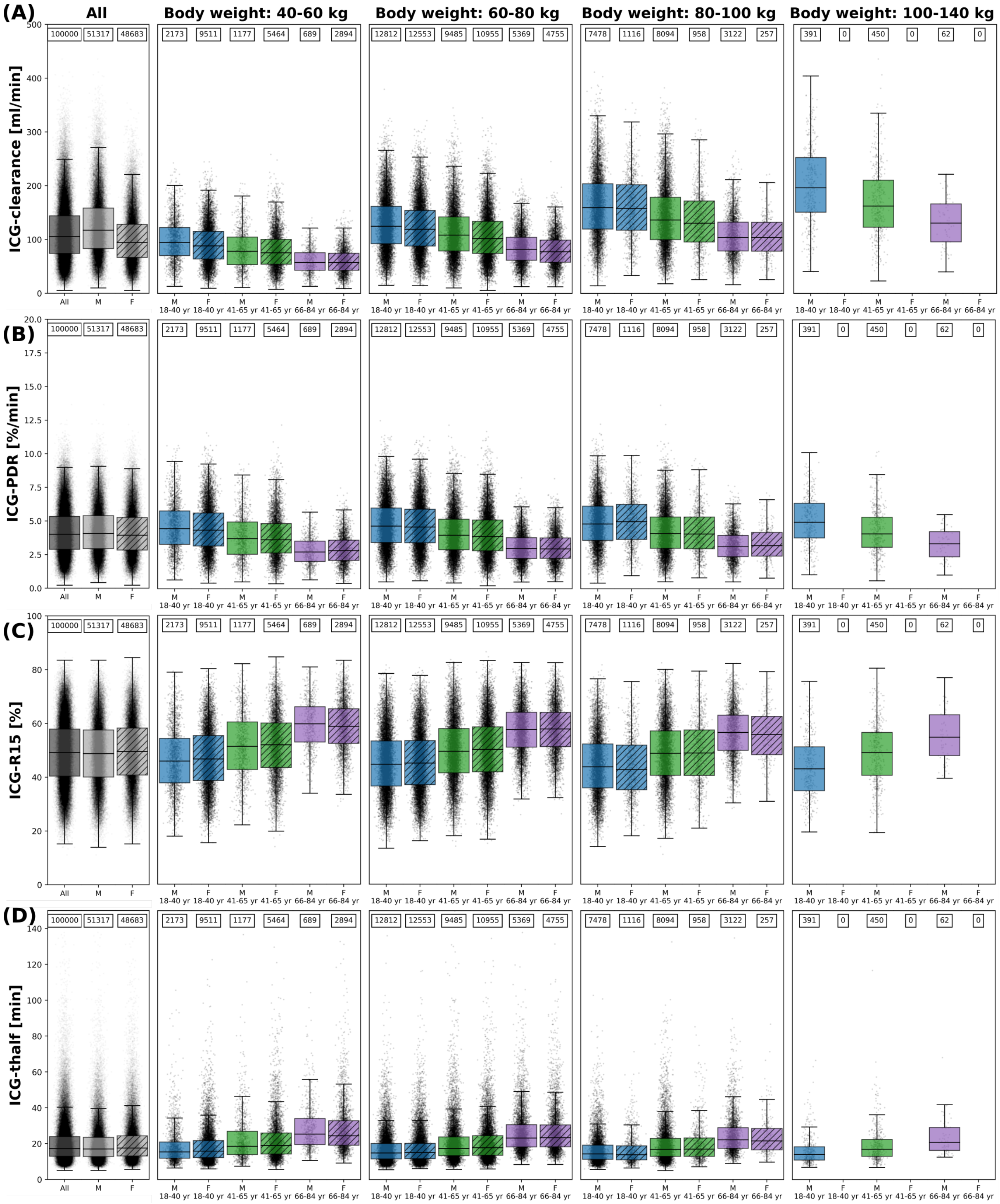


**Figure S3. ICG parameters *in silico* population with severe cirrhosis:** The dependency of **A** ICG-clearance, **B** ICG-PDR, **C** ICG-R15, and **D** ICG-$t_{1/2}$ on body weight, sex and age in n=100000 individuals. Results are stratified by body weight class (40-60 kg, 60-80 kg, 80-100 kg, 100-140 kg), age group (blue - 18-40 yr; green - 41-65 yr; purple - 66-84 yr) and sex (M - unshaded; F - shaded). No women exist in the data set with body weight $>$ 100 kg. The sample size of the respective subgroups are depicted above each boxplot. The box extends from the lower to upper quartile values of the data, with a line at the median with whiskers as defined by Tukey. Individual data points are plotted for all subgroups. Subjects are simulated with severe cirrhosis.
